# Biogenesis of DNA-carrying extracellular vesicles by the dominant human gut methanogenic archaeon

**DOI:** 10.1101/2024.06.22.600173

**Authors:** Diana P. Baquero, Guillaume Borrel, Anastasia Gazi, Camille Martin-Gallausiaux, Virginija Cvirkaite-Krupovic, Pierre-Henri Commere, Nika Pende, Stéphane Tachon, Anna Sartori-Rupp, Thibaut Douché, Mariette Matondo, Simonetta Gribaldo, Mart Krupovic

## Abstract

Extracellular vesicles (EVs) are membrane-bound particles secreted by cells from all domains of life and implicated in a variety of important processes, from intercellular communication to pathogenesis. Here, we characterize EVs produced by the dominant human gut methanogen, *Methanobrevibacter smithii*, which, unlike most archaea, contains a peptidoglycan cell wall. Using quantitative proteomics, we show that *M. smithii* EVs are enriched in various proteins responsible for chromatin structure, including histones, and DNA repair. Consistently, the *M. smithii* EVs carry DNA, with fragments covering the entire cellular chromosome. Notably, the EVs are strongly enriched in extrachromosomal circular DNA (eccDNA) molecules which originate from excision of a 2.9-kb chromosomal fragment and a proviral genome. The eccDNA encodes two of the key methanogenesis enzymes and could boost their expression inside the cells through the gene dosage effect. Furthermore, four of the top ten most abundant EV proteins are implicated in methanogenesis. Cryo-electron tomography (Cryo-ET) suggests that EVs are formed by budding from the cell membrane and are trapped under the cell wall prior to liberation through local disruptions in the cell wall. Collectively, our results reveal parallels with EV biogenesis in bacteria and suggest that *M. smithii* EVs facilitate the export of both cellular and viral DNA as well as key metabolic proteins in the gut environment, with potential impact on methane production.

## INTRODUCTION

Extracellular vesicles (EVs) are membrane-bound particles secreted into the extracellular environment by cells from all walks of life. EVs range in size from 20 to 400 nm in diameter and bud off from cellular membranes (Deatherage and Cookson, 2012; Gill et al., 2019). Although EVs were initially thought to carry cellular waste products, represent cell debris or artifacts of lipid aggregation (Coelho and Casadevall, 2019), it is now recognized that EVs play multiple important roles in diverse biological processes, including horizontal gene transfer, transport of metabolites, cell-to-cell communication, biofilm formation, virulence and antiphage defense (Douanne et al., 2022; Kuehn and Kesty, 2005; Ofir and Sorek, 2017; S et al., 2013; Schooling and Beveridge, 2006; Toyofuku et al., 2019). EVs derived from humans can promote certain pathologies, including cancer, Alzheimer’s and Parkinson’s diseases (Blommer et al., 2023; Bodart-Santos et al., 2023; Buzas, 2023; Chang et al., 2021). Thus, understanding the role of EVs across domains of life emerges as an important area of research.

Most of the research on EVs thus far has focused on eukaryotes and bacteria. However, a growing body of evidence suggests that phylogenetically diverse archaea from various ecosystems also produce EVs (Liu et al., 2021b). In particular, EVs have been described in hyperthermophilic archaea of the orders Sulfolobales, Thermococcales and Methanococcales as well as halophilic archaea of the order Halobacteriales (Choi et al., 2015; Erdmann et al., 2017; Gaudin et al., 2013; Gorlas et al., 2015; Marguet et al., 2013; Mills et al., 2024; Soler et al., 2008; Thiroux et al., 2021). Although the impact of EVs on archaeal communities remains to be fully understood, it is becoming increasingly clear that they play an active role in various ecosystems. The EVs produced by a thermoacidophile *Saccharolobus islandicus* (order Sulfolobales), the first archaeal EVs to be characterized, were found to carry an antimicrobial protein named ‘sulfolobicin’, which selectively inhibits the growth of other Sulfolobales species (Prangishvili et al., 2000). In contrast, *Thermococcus prieurii,* a sulfur-reducing hyperthermophilic archaeon, secretes EVs packed with elemental sulfur, likely to prevent the accumulation of cytotoxic levels of sulfur inside the cells (Gorlas et al., 2015). Archaeal EVs might also play an important role in horizontal gene transfer and DNA encapsulation within EVs appears particularly critical in extreme geothermal and acidic environments. Indeed, EVs produced by *S. islandicus*, *Thermococcus* and *Halorubrum* species were shown to carry chromosomal and/or plasmid DNA (Choi et al., 2015; Erdmann et al., 2017; Gaudin et al., 2013; Gaudin et al., 2014; Liu et al., 2021a; Soler et al., 2008). For instance, EVs from *S. islandicus* REY15A could transfer the plasmid-borne *pyrEF* locus into a plasmid-free auxotrophic *S. islandicus* strain (Liu et al., 2021a). Finally, under nutrient limiting conditions, *S. islandicus* EVs were shown to serve as a carbon and nitrogen source, promoting microbial growth (Liu et al., 2021a).

The mechanisms of EV biogenesis remain poorly understood, but it is evident that the structure and composition of the cell envelope play a central role in this process. In most bacteria, the cytoplasmic membrane is encased by a rigid peptidoglycan layer, which presents a barrier for EV release. Two major mechanisms of EV production have been proposed. The first mechanism is common to both monoderm (gram-positive) and diderm (gram-negative) bacteria and occurs through explosive lysis caused by phage infection, induction of prophages or action of autolysins. Accordingly, EVs produced through explosive lysis route (E-type EVs) are enriched in peptidoglycan digesting enzymes (Toyofuku et al., 2023). The second mode of EV release occurs via membrane blebbing (B-type EVs). In diderm bacteria, in which the peptidoglycan is covered by an outer membrane, the EVs are normally produced by blebbing of the outer membrane. Recent super-resolution microscopy analysis of EV biogenesis showed that EVs of monoderm bacteria can be also produced by budding/blebbing from the cytoplasmic membrane (Jeong et al., 2022). These EVs undergo a ‘waiting’ period whereby they are trapped between the membrane and peptidoglycan until local openings in the peptidoglycan layer allow their release (Jeong et al., 2022). EVs produced via explosive lysis are larger and more variable in diameter compared to the EVs produced by blebbing from the cytoplasmic membrane (Jeong et al., 2022; Jiang et al., 2024; Toyofuku et al., 2023).

The envelope of most archaeal cells consists of a cytoplasmic membrane covered by a paracrystalline protein surface (S-) layer (Albers and Meyer, 2011). Similarly, all archaeal EVs characterized thus far are covered by the cellular S-layer, consistent with budding from the cytoplasmic membrane (Bonanno et al., 2019; Liu et al., 2021a; Prangishvili et al., 2000). However, some archaea, most notably, methanogenic archaea of the order Methanobacteriales, the dominant group of archaea in the animal gastrointestinal tract (GIT) (Thomas et al., 2022), have a peptidoglycan polymer surrounding the cell membrane (Albers and Meyer, 2011; Meyer and Albers, 2020; Steenbakkers et al., 2006). Whether gut methanogens with a rigid cell wall can produce EVs remains unknown. This question is of particular interest given that EVs produced by gut bacteria play an important role in regulating the intestinal microenvironment, modulating both intermicrobial and microbe-host interactions (Liang et al., 2022; Wang et al., 2024).

Here, we characterize the composition and biogenesis of EVs produced by *Methanobrevibacter smithii*, the dominant archaeon in the human GIT, accounting for over 90% of the gut archaeome (Borrel et al., 2020; Chibani et al., 2022; Dridi et al., 2009; Eckburg et al., 2005; Mihajlovski et al., 2010). *M. smithii* is a strict anaerobe which obtains energy by reducing carbon dioxide into methane using molecular hydrogen as electron donor (Miller et al., 1982). *M. smithii* cells have an ovococcoid morphology and are surrounded by a peptidoglycan polymer as a primary cell wall component. We demonstrate that *M. smithii* EVs carry fragments of chromosomal DNA, are enriched in proviral DNA and extrachromosomal circular DNA molecules encoding key proteins of methanogenesis. Consistently, the EV proteome is dominated by proteins involved in DNA metabolism and includes several proteins implicated in methanogenesis. Furthermore, cryo-electron tomography (cryo-ET) analysis shows that *M. smithii* EVs accumulate in the pseudo-periplasmic space before being released into the extracellular environment through local disruptions in the *M. smithii* cell wall, revealing parallels with the mechanism of EV biogenesis by monoderm bacteria.

## RESULTS

### Characterization of EVs from *Methanobrevibacter smithii*

To investigate the potential production of EVs in members of the order Methanobacteriales, we chose the type strain *Methanobrevibacter smithii* PS (ATCC 35061) as a model organism. We established a protocol for obtaining highly purified EV preparations from *M. smithii*, a prerequisite for biochemical characterization **(Fig. 1A)**. In particular, the EVs were isolated from exponentially growing *M. smithii* cultures to limit the contamination of the EV preparations with cell debris. All sub-cellular particles from filtered cell-free supernatants were collected by ultracentrifugation and subsequently loaded at the bottom of an iodixanol (OptiPrep™) floatation gradient in which particles containing lipids are expected to float up due to their lower density compared to proteinaceous particles (**Fig. 1A**). The gradient was fractionated, particles in the collected fractions washed and concentrated in phosphate buffered saline (PBS) solution (**Fig. 1A**). The SDS-PAGE (**Fig. 1B**) and transmission electron microscopy (TEM; **Fig. 1C**) analyses showed that the EVs of variable diameters were predominantly present in the gradient fractions with the densities between 1.11 and 1.13 g/mL.

**Figure 1.**
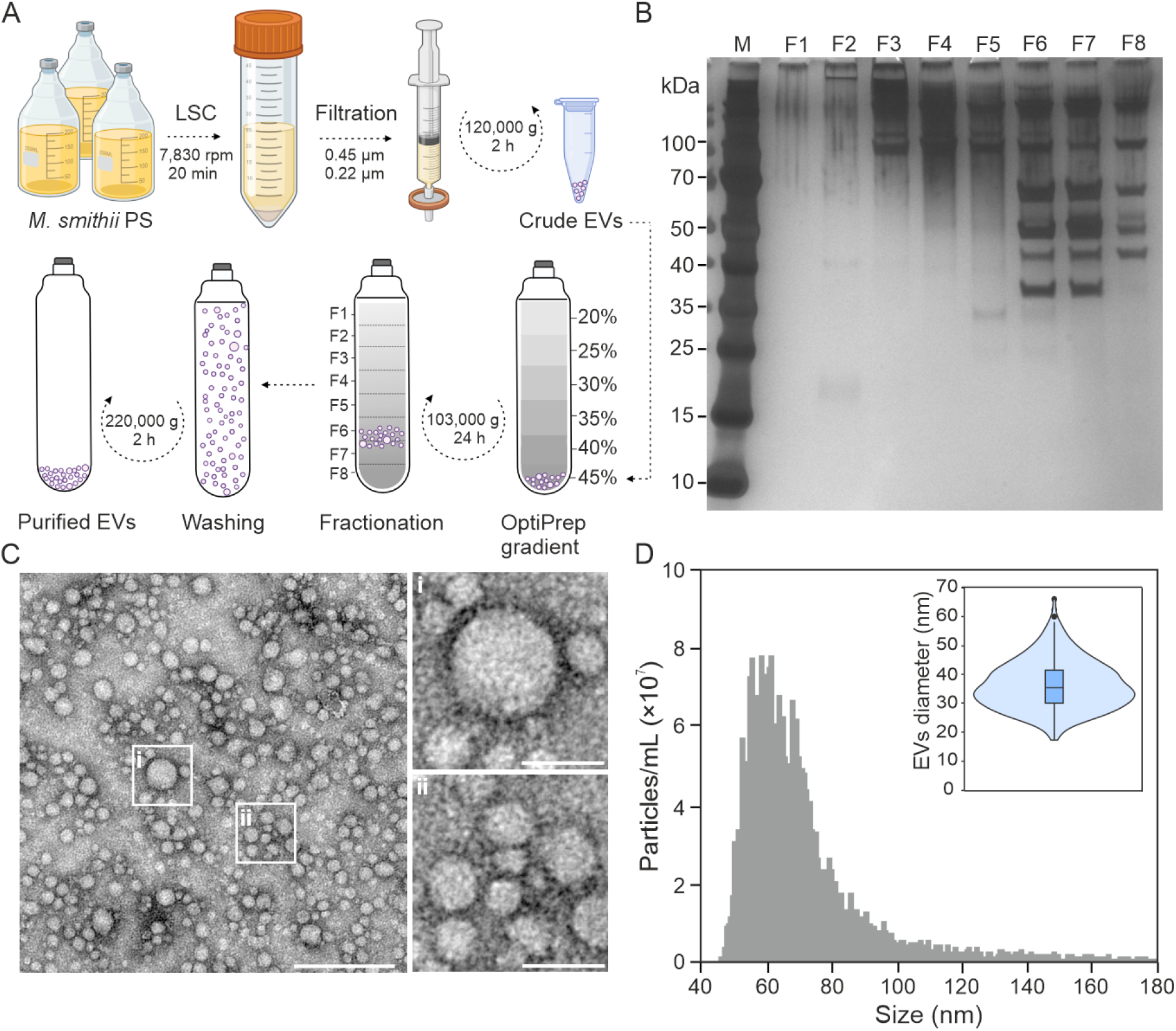
Characterization of EVs produced by the methanogenic archaeon *Methanobrevibacter smithii* PS. **A**, Purification protocol established for *M. smithii* EVs. Crude EVs were obtained by ultracentrifugation of cell-free supernatants from *M. smithii* cultures. The pelleted crude EVs were resuspended in 45% OptiPrep™, and layers of lower concentrations were loaded on top. After ultracentrifugation of the OptiPrep gradient, fractions containing EVs were collected and combined. The collected EVs were washed in 10 times the volume of PBS solution and collected by ultracentrifugation. The pelleted purified EVs were then resuspended in PBS buffer. Panel created with BioRender.com. **B**, SDS-PAGE gel of fractions collected after OptiPrep gradient. Fractions 6 and 7 contain the largest number of EVs. **C**, Transmission electron micrographs of *M. smithii* EVs. The main image displays EVs of different sizes. Scale bar, 200 nm. The upper right panel depicts a vesicle of approximately 60 nm in diameter. The bottom right panel displays smaller EVs with sizes ranging from 18-30 nm. Scale bar for both right panels: 50 nm. Samples were negatively stained with 2% uranyl acetate. **D**, Particle size distribution determined by NanoFCM. The violin plot displays the relative EV diameter when calculated manually (n=412). The width of the distribution indicates the frequency of occurrence.

The surface of negatively stained *M. smithii* EVs showed no apparent presence of the cell wall components (**Fig. 1C**). The size distribution of purified EVs was estimated using NanoFCM (Nano FCM Inc., Xiamen, China), a flow cytometer specifically designed for the analysis of nano-sized particles such as EVs and viruses. The majority of particles measured using NanoFCM had diameters ranging from 45 to 100 nm (median diameter of 65 nm; **Fig. 1D**). Given that the detection limit of NanoFCM is ∼40 nm, we also determined the size of *M. smithii* EVs by measuring the relative diameters of 412 particles imaged by TEM. The manually measured median diameter of EVs was 32 nm (min=17 nm, max=67 nm), indicating that NanoFCM measurements did not detect the majority of EVs produced by *M. smithii* (**Fig. 1D**). No particles larger than 67 nm in diameter were observed by TEM, likely due to their lower abundance compared to the small-diameter EVs (**Fig. 1C**). Collectively, our results suggest that *M. smithii* EVs are spherical, display a wide variation in diameter (∼20-100 nm), with the majority of particles having a diameter of ∼30 nm.

### *M. smithii* EVs preferentially enclose a 2.9 kb circular DNA molecule

EVs produced by hyperthermophilic and halophilic archaea were previously shown to carry DNA (Erdmann et al., 2017; Gaudin et al., 2013; Liu et al., 2021a; Soler et al., 2008). To determine whether EVs from *M. smithii* also carry genetic material, the purified EVs were treated with DNase I to eliminate any extracellular DNA, followed by DNA extraction. Sequencing of the DNA purified from the EVs using the Illumina platform produced reads that mapped to the entire *M. smithii* chromosome with an average sequencing depth of 15× (**Fig. 2A**, **Fig. 2B ii**). The small size of the *M. smithii* EVs precludes the encapsulation of a continuous complete chromosome within distinct EVs. Instead, it is likely that overlapping genomic fragments of varying sizes are randomly enclosed within the EVs, as has been reported for EVs from hyperthermophilic archaea (Gaudin et al., 2013; Liu et al., 2021a; Soler et al., 2008) as well as bacterial EVs (Biller et al., 2017).

**Figure 2.**
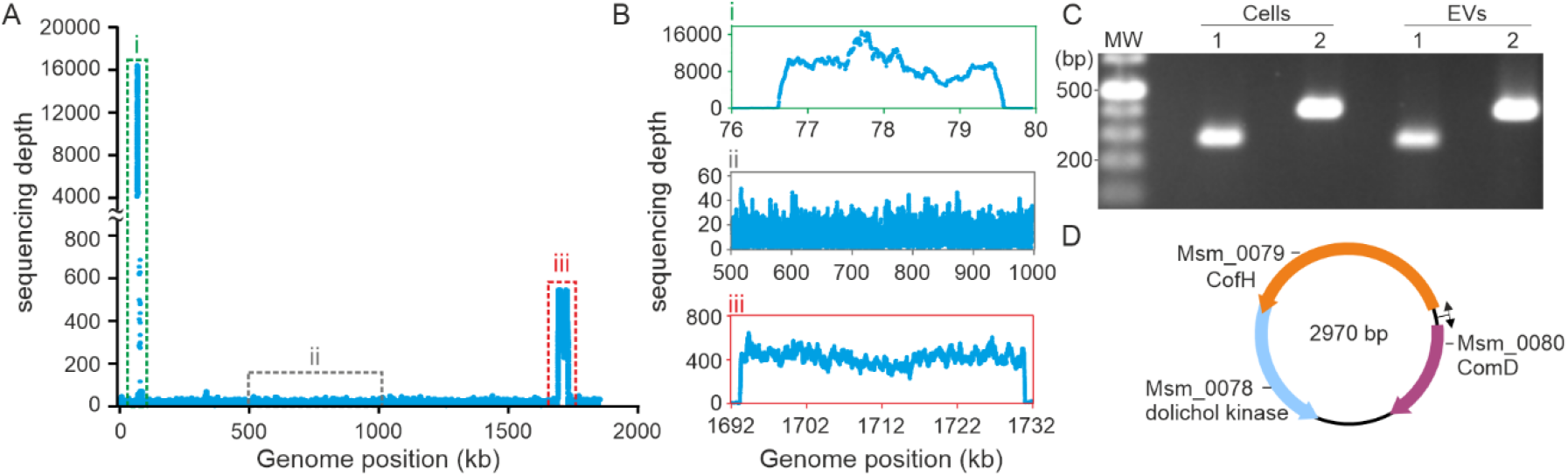
DNA content of *M. smithii* EVs. **A**, Coverage depth across the *M. smithii* PS chromosome. Each dot represents the coverage at the indicated position. **B**, Coverage depth across distinct regions of the *M. smithii* PS chromosome: region i (nucleotide coordinates 76-80 kb) giving rise to a circular element of 2.9 kb; region ii (nucleotide coordinates 500-1000 kb), representing the typical sequencing depth across the genome; region iii (nucleotide coordinates 1692 – 1732 kb), encompassing the MSTV1 provirus. **C**, Detection of the integrated and excised form of the 2.9 kb DNA fragment in both cell and purified EV preparations. The agarose gel electrophoresis shows the PCR amplified products: lane 1, integrated element within the host chromosome (expected size: 291 bp); lane 2, excised and circularized form of the 2.9 kb DNA element (expected size: 433 bp). **D**, Genome map of the circular element. The ORFs are represented by arrows indicating the direction of transcription. CofH, 5-amino-6-(D-ribitylamino)uracil:L-tyrosine 4-hydroxybenzyltransferase; ComD, sulfopyruvate decarboxylase subunit alpha.

Unexpectedly, two chromosomal regions displayed a remarkably high sequencing depth (**Fig. 2A**, **Fig. 2B i and iii**). The first, with an average sequencing depth of 9106×, i.e., ∼600× higher than the overall average, spans a chromosomal region between nucleotide positions 76,619 and 79,634 (**Fig. 2B i**). The reads covering this region were assembled into a circular contig of 2970 bp, suggesting the excision and circularization of the corresponding chromosomal region. To verify this assumption, we designed primers amplifying either the chromosomal locus or the excised element. Polymerase chain reaction (PCR) analyses and subsequent sequencing confirmed the presence of “integrated” and “excised” forms in both cells and EVs (**Fig. 2C**, **Fig. S2-S4**), indicating that this locus indeed undergoes excision and circularization.

The circular element carries three genes (Msm_0078–Msm_0080) none of which encode functions typical of mobile genetic elements (MGEs), i.e., integrases, transposases, or genome replication proteins. Instead, the three genes are implicated in different metabolic pathways (**Fig. 2D)**. Msm_0078 encodes a putative dolichol kinase (UniRef: A5UJA5), an enzyme involved in the synthesis of CDP-diglyceride, a compound that plays a key role in the biosynthesis of phosphoglycerides, one of the main structural components of biological membranes (Gaillard et al., 1983). The two other genes, Msm_0079 and Msm_0080, encode 5-amino-6-(D-ribitylamino)uracil:L-tyrosine 4-hydroxybenzyltransferase (CofH; A5UJA6) and sulfopyruvate decarboxylase subunit alpha (ComD; A5UJA7), respectively. Whereas CofH catalyzes the production of 7,8-didemethyl-8-hydroxy-5-deazariboflavin, the precursor of the redox coenzyme F_420_, ComD is implicated in coenzyme M (CoM) biosynthesis (Graham, 2011), both playing a critical role in methanogenesis. In CO_2_-reducing methanogens, cofactor F_420_ is a crucial electron transporter, providing electron from H_2_ to reduce the methenyl group into methyl. In all types of methanogenesis pathways, CoM is the terminal methyl carrier before the formation of methane by the methyl-coenzyme M reductase complex.

Notably, the borders of excision of the circular element fall within protein coding genes, truncating genes Msm_0077 and Msm_0081 coding for thymidylate kinase (A5UJA4) and sulfopyruvate decarboxylase subunit beta (A5UJA8), respectively (**Fig. S2**). The potential functions of such circular molecules in *M. smithii* cells are intriguing. Given the presence of putative promoters in front of the two genes involved in methanogenesis (Msm_0079 and Msm_0080), the additional gene copies from eccDNA must boost the expression of the corresponding genes and have an impact on methanogenesis.

### *M. smithii* EVs carry viral DNA

The second chromosomal region with a high sequencing depth of 420× spans 38,824 bp (genome positions 1,693,231-1,732,055; **Fig. 2A**, **Fig. 2B iii**) and corresponds to a previously reported provirus in the genome of *M. smithii* PS (Krupovic et al., 2010). We recently demonstrated that this provirus, dubbed MSTV1, is sporadically reactivated in a small fraction of the *M. smithii* population, producing extracellular virus particles with a siphovirus-like morphology (Baquero et al., under review). MSTV1 is present in 20% of all sequenced *M. smithii* strains and is likely to be the most abundant archaeal virus in the human gut (Baquero et al., under review). PCR analyses revealed the excised form of the virus genome in both *M. smithii* cells and EVs (**Fig. S5**). The excision takes place at the proviral attachment sites and hence is mediated by the virus-encoded site-specific integrase of the tyrosine recombinase superfamily. The high sequencing depth of the provirus compared to the flanking regions (420× versus 15×) suggests that this locus in EVs is represented not only by the randomly incorporated fragments of the chromosomal DNA (as the rest of the chromosome) but also the actively excised viral DNA. Thus, the larger EVs observed by nanoFCM (**Fig. 1D**) may carry the complete viral genome and facilitate virus spread to non-infected cells in the population. This hypothesis could not be confirmed experimentally using the *M. smithii* strains available in the laboratory, either due to inability of the EVs to overcome the cell wall barrier of the target cells or due to resistance mechanisms that remain to be understood.

### *M. smithii* EVs are enriched in DNA-binding and DNA repair proteins

To assess the protein composition of the *M. smithii* EVs, we performed quantitative proteomics analysis of the *M. smithii* cells and EVs using mass spectrometry. The analyses led to the identification of 1073 proteins in cells and 417 proteins in EVs which were present in all three biological replicates (**Fig. S6**). None of the proteins were exclusive to the EVs **(Table S1)**. The number of proteins found in *M. smithii* EVs is similar to that described in EVs produced by other archaea, such as the hyperthermophilic archaeon *Saccharolobus islandicus* (413 proteins) and the halophilic archaeon *Halorubrum lacosprofundi* (447 proteins) (Erdmann et al., 2017; Liu et al., 2021a).

Archaeal Clusters of Orthologous Groups (arCOG) were used to classify the potential functions of the *M. smithii* EV proteins (Makarova et al., 2015) **(Table S2)**. Proteins from the arCOG category P (Inorganic ion transport and metabolism) were found exclusively in the cellular proteome (**Fig. 3A**). In contrast, EVs were more abundant in proteins of the arCOG category J (Translation, ribosomal structure, and biogenesis), comprising 19% of the total EV proteins, compared to 13% in the *M. smithii* cell proteome. EVs also exhibited enrichment in proteins from the arCOG categories C (Energy production and conversion), E (Amino acid transport and metabolism), F (Nucleotide transport and metabolism), G (Carbohydrate transport and metabolism), and O (Posttranslational modification, protein turnover, chaperones) (**Fig. 3A)**. The presence of proteins from nearly all arCOG categories in the EVs suggests that most of the proteins are incorporated non-selectively, likely by entrapment of the cytosolic and membrane contents, as previously suggested for other EVs (Liu et al., 2021a). Indeed, there is a strong positive correlation between the abundance of the proteins in the cells and their abundance in EVs (Pearson correlation coefficient r=0.9189; **Fig. 3B**).

**Figure 3.**
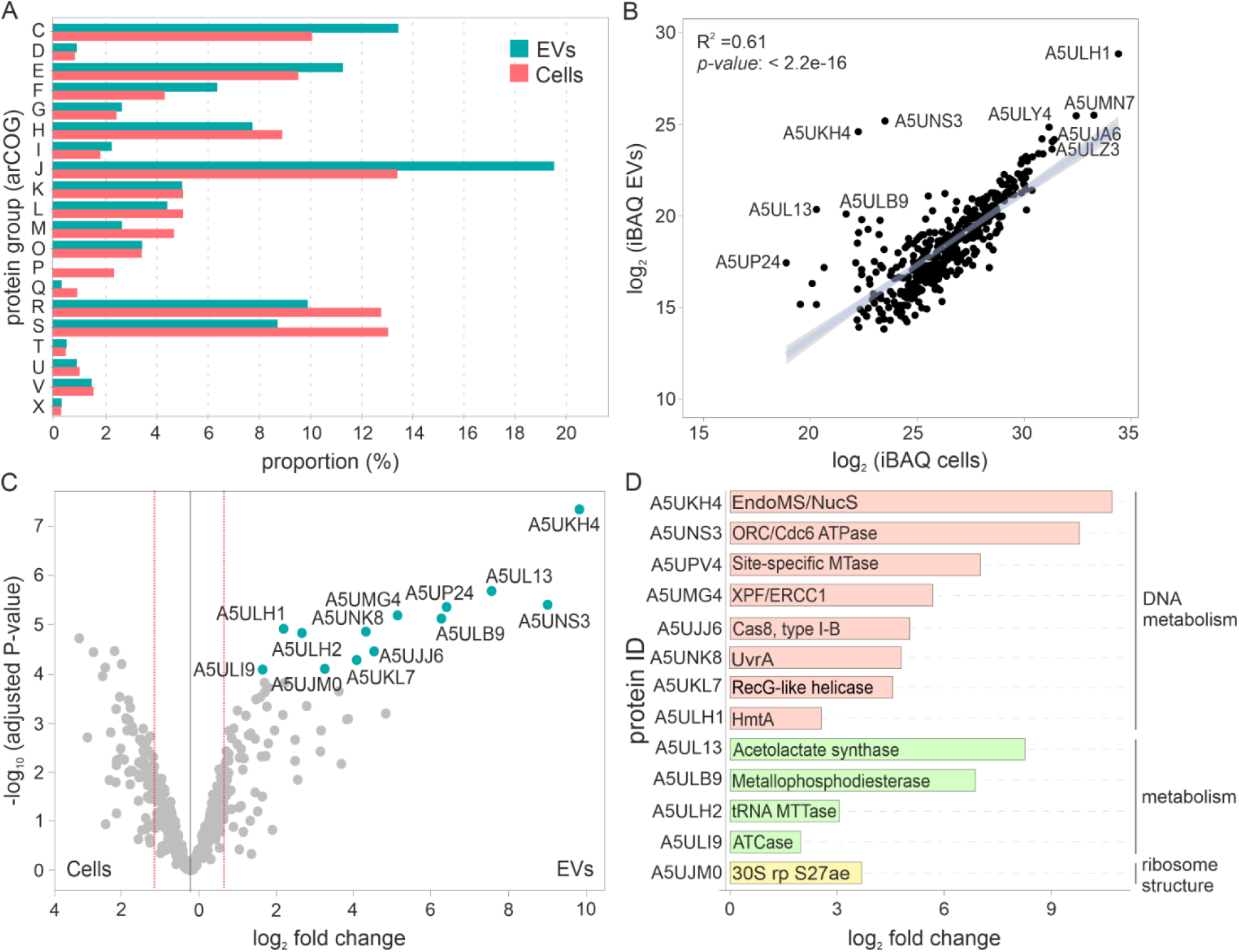
Analysis of protein content of *M. smithii* EVs. **A**, Functional classification of proteins identified in purified *M. smithii* EVs using archaeal clusters of orthologous groups (arCOGs). Annotations for the arCOG categories are provided in **Table S2**. **B,** Correlation between protein abundances in *M. smithii* EVs and cells. **C**, Volcano plot comparing protein content of *M. smithii* EVs and cells. Read lines highlight the threshold for enrichment t (p = 0.05 and |fold change| = 1.5). Differential protein abundances and adjusted p-values were calculated using the package DEP (Differential Enrichment analysis of Proteomics data). **D**, Proteins significantly enriched in EVs when compared to proteins in cellular fraction.

Label-free intensity-based absolute quantification (iBAQ) of the *M. smithii* EV proteome showed that the three most abundant proteins (**Table S3**) are the three histone paralogs encoded by *M. smithii* PS (A5ULH1, A5UJP0, A5UMN7) (Stevens et al., 2020). Histone-fold proteins have been shown to mediate genome compaction in the order Methanobacteriales (Hocher et al., 2022; Rojec et al., 2019; Stevens and Warnecke, 2023). These observations suggest that the DNA in the EVs is in the form of DNA-histone complexes. Notably, the top 4 and top 6 overall most abundant proteins in the EVs are also associated with DNA metabolism (**Table S3**). The top 4 most abundant protein is a homolog of the Orc1/Cdc6 AAA+ ATPase (A5UNS3) encoded by the MSTV1 provirus (**Fig. 2B, Table S3**) and predicted to be involved in a regulatory circuit controlling the switch between the temperate and lytic state of MSTV1 (Baquero et al. under review). Notably, Orc1/Cdc6 is the only provirus protein found in all three replicates of the EVs proteome, confirming that the viral DNA detected in the EVs does not originate from contaminating virus particles. This result is consistent with EV purification strategy used, where proteinaceous virus particles are not expected to co-float with the lipid-containing membrane vesicles. The top 6 most abundant protein in the EVs is the endonuclease EndoMS/NucS (A5UKH4) (**Table S3**), a multifunctional enzyme involved in DNA repair processes such as nucleotide excision repair, mismatch repair, and deaminated base repair (Ishino et al., 2016; Zhang et al., 2020). Presumably, EndoMS/NucS is incorporated into EVs along with its damaged DNA substrates.

Notably, four of the top 10 most abundant EV proteins are related to methanogenesis (**Table S3**). In particular, N^5^-methyl-tetrahydromethanopterin:coenzyme M methyltransferase subunit H (MtrH), the above mentioned 5-amino-6-(D-ribitylamino)uracil:L-tyrosine 4-hydroxyphenyl transferase CofH involved in F_420_ biosynthesis, methyl-coenzyme M reductase subunit gamma and F_420_-dependent methylenetetrahydromethanopterin dehydrogenase (A5ULY4, A5UJA6, A5ULZ3, and A5UMI1, respectively). CofH is the fourth most abundant protein in the total *M. smithii* proteome (**Table S4**). It is tempting to speculate that the high expression of this protein could be at least partly caused by the extra gene copies borne on the excised circular DNA molecules.

Another notable protein present in high abundance in the EVs (top 14) is an adhesin (Msm_1398; A5UN25) with an N-terminal pectin lyase domain and 4 tandem immunoglobulin (Ig)-like domains at the C-terminus (**Table S3**). Adhesins were suggested to play a key role in ensuring the persistence of *M. smithii* PS in the distal intestine (Samuel et al., 2007). Similarly, Ig-like domains are specifically enriched in gut-associated archaeal viruses and are thought to mediate adhesion of virus particles to various substrates, including cell surface exposed glycans and the eukaryotic mucus layer (Medvedeva et al., 2023). The adhesins present in EVs could play a similar role. Notably, the same protein is also one of the most abundant adhesins in the total *M. smithii* proteome (top 11; **Table S4**).

Next, we computed the differential protein abundance in the EVs compared to the cellular proteome (log2 > 1.5, adjusted *P*-value < 0.05) (**Fig. 3C**). This analysis identified 13 proteins that are significantly enriched in EVs compared to the total cell proteome (**Table S5**). Besides the endonuclease EndoMS/NucS (A5UKH4), one of the histone paralogs (A5ULH1), and the viral Orc1/Cdc6 AAA+ ATPase (A5UNS3), several other DNA metabolism and repair proteins were found to be among the most significantly enriched proteins in *M. smithii* EVs when compared to the cell proteome (**Figs. 3B-D**). These include the UvrABC nucleotide excision repair system subunit UvrA (A5UNK8) (Crowley et al., 2006), nucleotide excision repair XPF/ERCC1 family helicase-nuclease (A5UMG4), RecG-like ATP-dependent DNA helicase (A5UKL7), adenine-specific DNA methyltransferase (A5UP24) and type I-B CRISPR-associated protein Cas8 (A5UJJ6) (Cass et al., 2015). The high abundance of proteins related to chromatin organization and DNA metabolism in the EVs suggests that the DNA fragments covering the entire *M. smithii* chromosome could be generated during the DNA repair processes, in particular, nucleotide excision repair pathway, which generates fragments of damaged DNA (Marshall and Santangelo, 2020).

### Cryo-ET provides insights into EV production in *M. smithii* cells

To gain insights into the biogenesis of *M. smithii* EVs, we analyzed the exponentially growing *M. smithii* cells by cryo-electron tomography (cryo-ET) (Baumeister, 2022). Spherical EV-like particles 12-45 nm in diameter (n=31) were visualized in the cells, usually trapped in ‘pockets’ between the cytoplasmic membrane and the archaeal peptidoglycan polymer (periplasmic space) (**Fig. 4A, Video S1**). Multiple EVs with different diameters could be observed in a single cell. The reconstructed tomograms also revealed potential extracellular EVs in the close proximity of the imaged cells. Consistent with our TEM and proteomics analyses of the purified EVs, both the periplasmic and extracellular EVs lack an apparent peptidoglycan coat.

**Figure 4.**
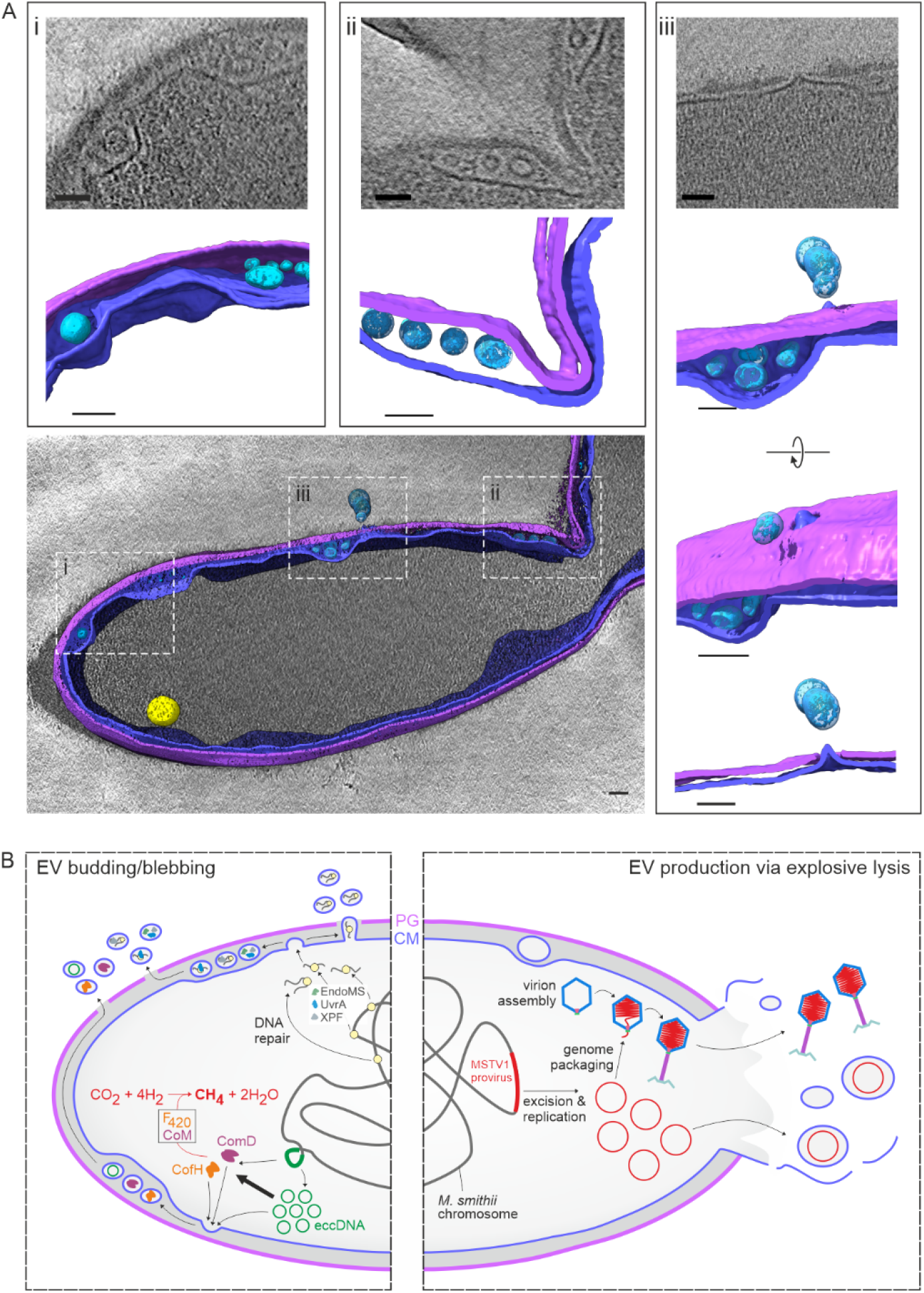
Biogenesis of extracellular vesicles (EVs) by *M. smithii* cells. **A,** Tomographic reconstruction of a *M. smithii* cell producing EVs. The central panel shows a segmented, surface-rendering display of the reconstructed tomogram of a *M. smithii* cell producing EVs. Panels i and ii exhibit EVs trapped in ‘pockets’ between the cytoplasmic membrane and the archaeal peptidoglycan. Panel iii displays different tilts of a region in the cell where the peptidoglycan is locally disrupted, with the opening coinciding with the protrusion of the cell membrane and the presence of EVs outside of the cell. The reconstruction displays the following cellular components: membrane (purple blue), cell wall (purple), EVs (light blue), storage granule-like structure (yellow). Scale bars, 50 nm. **B,** A schematic representation of EVs biogenesis and release by a *M. smithii* cell. DNA fragments from random chromosomal loci are incorporated into the *M. smithii* EVs. Two chromosomal regions, corresponding to the MSTV1 provirus and a 2.9-kb extrachromosomal circular DNA (eccDNA) molecule, are enriched in the *M. smithii* EVs. The eccDNA molecules encode two key methanogenesis enzymes that could boost methanogenesis through a gene dosage effect. Consistent with the presence of DNA, proteins responsible for chromatin structure and DNA repair are also enriched in *M. smithii* EVs. Our cryo-ET results show that small EVs of approximately 12-45 nm in diameter are trapped in ‘pockets’ between the cytoplasmic membrane (CM) and the peptidoglycan (PG). We hypothesize that the release of small (∼32 nm) and large (>100 nm) EVs in *M. smithii* occurs via distinct routes, budding/blebbing (left) and explosive virus-mediated lysis (right), respectively.

Our data provides possible clues on how the EVs pass through the peptidoglycan polymer, a question which is yet to be addressed also for bacterial EVs. We observed an area in the vesiculating cell that corresponds to a local opening (22 nm in diameter) in the cell wall, which colocalizes with the outward protrusion of the cell membrane and the presence of EVs outside of the cell (**Fig. 4A**). While it seems plausible that *M. smithii* EVs traverse through these areas, the mechanism for archaeal cell wall degradation and the components involved in this process remain unclear. We note that the cell which was captured in the process of EV biogenesis is undergoing division (**Fig. 4A**), a time point in the cell cycle during which the cell wall is subjected to active remodeling. In monoderm bacteria, which encompass a thick cell wall of 20-40 nm, the action of phage- or host-encoded peptidoglycan-degrading enzymes has been demonstrated to facilitate the transit across the cell wall (Bose et al., 2020; Lee et al., 2009; Resch et al., 2016; Toyofuku et al., 2017). Consistently, the bacterial EVs typically carry either phage-encoded lysins or cellular peptidoglycan digesting autolysins (Hayashi et al., 2002; Liu et al., 2022; Wang et al., 2018). Notably, the PeiW-like endolysin (Peptidase_C71 family) encoded by MSTV1 (Medvedeva et al., 2023) or other *M. smithii* lysins were not detected in EVs by proteomics analysis.

## DISCUSSION

Here, we characterized the EVs from *Methanobrevibacter smithii*, the dominant methanogen in the human gut with a peptidoglycan cell wall. We show that *M. smithii* EVs carry three types of DNA molecules which vastly differ in their abundancies. These correspond to (i) random genomic loci distributed across the *M. smithii* chromosome, (ii) MSTV1 provirus and (iii) an extrachromosomal circular DNA (eccDNA) molecule of 2.9 kb. The frequency of incorporation into EVs likely depends on the availability (copy number) and dimensions of the corresponding molecules in the cell during the EV biogenesis. The fragments cumulatively covering the entire chromosome could be generated during various DNA repair processes, including DNA mismatch and nucleotide excision repair (NER) pathways. Indeed, EndoMS/NucS, the top 6 most abundant protein in EVs, is the major player in mismatch repair in most archaeal lineages (Ahmad et al., 2020; Ishino et al., 2016; Ishino et al., 2018). By contrast, UvrA and XPF proteins, which are significantly enriched in EVs, are key players in the NER pathway, with UvrA being responsible for recognition of DNA helix distorting lesions and XPF implicated in nicking the damaged DNA strand (Marshall and Santangelo, 2020). In bacteria, two different nucleases, XPF and XPG, nick the damaged DNA downstream and upstream of the lesion, respectively. However, since archaea lack homologs of XPG, it was suggested that XPF could cut the DNA on both sides of the lesion (Marshall and Santangelo, 2020). Alternatively, it has been suggested that EndoMS/NucS could participate in the NER pathway together with XPF (Zhang et al., 2020). Thus, small linear DNA fragments generated through the NER pathway could be enclosed within the EVs. Due to the inherently random distribution of mutations in the population, different cells would be expected to produce EVs with random genomic fragments. Notably, the two high-frequency regions (the provirus and eccDNA) are also incorporated into EVs as part of the chromosomal DNA (i.e., in their integrated form), presumably with the same frequency as the flanking regions, as confirmed by the PCR analysis with the primers amplifying across the integration borders. However, the presence of the provirus and eccDNA in the extrachromosomal form increases their copy number and hence the probability of being incorporated into EVs, partly explaining their higher abundance in the EVs.

The cell envelope architecture of the *M. smithii* cells could be another important factor underlying the incorporation frequency of different cargo DNA molecules. The size of the EVs is likely to be dictated, at least in part, by the diameter of the (non-lethal) openings in the peptidoglycan cell wall. Accordingly, EVs with diameters small enough to pass through the peptidoglycan holes can be expected to enclose only DNA molecules of relatively small size. The median diameter of the *M. smithii* EVs measure from TEM images was ∼32 nm, consistent with the EVs observed by cryo-ET. Note that the diameter of the MSTV1 capsid in which the 38-kb viral genome is packaged under pressure is ∼65 nm (Baquero et al., under review). Thus, although histone proteins detected in the EVs are likely to condense the cargo DNA, the volume and hence the diameter of the EVs enclosing the viral DNA is still likely to exceed 65 nm. NanoFCM analysis revealed that a minor fraction of the EVs have diameters between 100 and 180 nm, which could be sufficient for carrying the full viral genome. Although it is conceivable that the EVs could be hijacked by MSTV1 to provide an alternative route for virus spread in the population, it is also possible that viral genomes are excreted from the cells as a form of an innate antiviral defense strategy. Notably, one of the most abundant proteins in the EVs corresponds to the viral Orc1/Cdc6 AAA+ ATPase (A5UNS3), an enzyme thought to be involved in the switch from the temperate to the lytic state of MSTV1 (Baquero et al., under review). Export of this protein from the cell could ensure stable lysogeny, thereby ensuring the cell’s survival. The presence of viral DNA in EVs has been previously reported in the hyperthermophilic archaeon *Thermococcus nautili* (Gaudin et al., 2014). Prophage genomes were also reported in EVs from *Vibrio cholerae* (Langlete et al., 2019). In eukaryotes, EVs have been shown to incorporate and transport infectious viral particles and viral genomes, such as rotaviruses, enteroviruses or noroviruses, to recipient cells (Chen et al., 2015; Kerviel et al., 2021; Santiana et al., 2018). Our findings further underscore the complex interplay between viruses and EVs.

Random chromosomal fragments, plasmids and viral DNA were previously detected in EVs produced by halophilic and hyperthermophilic archaea (Erdmann et al., 2017; Gaudin et al., 2014; Liu et al., 2021a; Soler et al., 2008). By contrast, the small eccDNA molecules encoding metabolic genes described here were previously not reported in archaea. These molecules resemble the eccDNA described in diverse eukaryotes, including plants, nematodes, ciliates, yeast, and mammals (Yang et al., 2022). eccDNA have been known in eukaryotes for over five decades, yet their significance and mechanisms of biogenesis remain enigmatic. Different strategies for the formation of circular extrachromosomal elements have been proposed, such as spurious homologous recombination between tandem copies (Shibata et al., 2012), DNA damage repair processes or genome arrangements caused by transposable elements (Zhang et al., 2023). Additionally, polymerase slippage in regions with repetitive sequences can create DNA loops during replication or repair that, when excised, result in the extrachromosomal circular elements (Dillon et al., 2015; Yang et al., 2022). We hypothesize that the 2.9 kb circular DNA element found both in *M. smithii* cells and EVs is excised from the chromosome by a similar, currently not understood mechanism. Regardless, when inside the cell, the eccDNA is bound to increase the copy number of genes it carries, thereby boosting their expression. In the case of the *M. smithii* eccDNA, two of the three genes encode key enzymes involved in methanogenesis, with one of these proteins being the fourth and eight most abundant protein in the *M. smithii* and EV proteomes, respectively. In addition, three other methanogenesis-related enzymes are among the top 10 most abundant EV proteins. Thus, EVs could provide the means of efficient and rapid discharge of the extra gene copies and proteins once they are no longer needed in high quantities.

In hyperthermophilic archaea, EVs provide means for horizontal gene transfer. However, given that Methanobacteriales cells are surrounded by a rigid cell wall, fusion of EVs with the cytoplasmic membrane might not be straightforward, if at all possible. It is thus more likely that EVs provide means for the discharge of damaged or viral DNA, or surplus components, both proteins and genes, that outlived their role under given conditions. Notably, the liberation of EVs trapped in the periplasmic space might not be the only possible outcome. Conceivably, the EVs could fuse back to the cytoplasmic membrane, reintroducing their cargo (e.g., eccDNA) into the cell. Under this scenario, the *M. smithii* EVs would serve an important regulatory role, finetuning methanogenesis to changing environmental or cellular cues.

The mechanism of EV biogenesis in prokaryotes is an outstanding question. Our cryo-ET analysis shows that the small-diameter *M. smithii* EVs are present in the pseudo-periplasmic space prior to their release into the extracellular milieu. This observation is inconsistent with blebbing through the holes in the peptidoglycan layer due to high turgor pressure, one of the models suggested for some bacterial EVs (Brown et al., 2015). Instead, the process appears to be similar to that described in a recent super-resolution microscopy analysis of EV biogenesis in *Staphylococcus* species, whereby EVs undergo a ‘waiting’ period in the periplasm until local openings in the peptidoglycan layer allow their release (**Fig. 4B**) (Jeong et al., 2022). Bacterial EVs carrying viral genomes are thought to be generated during explosive virus-mediated lysis. It is possible that the MSTV1 genome containing *M. smithii* EVs are also generated through a similar process (**Fig. 4B**). During the lytic virus cycle, a fraction of the viral genomes would be encapsidated into virions using the viral genome packaging ATPase, whereas the genomes which were not packaged at the time of lysis could be released into the environment within larger EVs. Indeed, large openings in the peptidoglycan cell wall required for the release of the viral genome-carrying EVs could have detrimental effects on the cell integrity. Thus, the mechanisms generating the small (∼32 nm) and larger (>100 nm) EVs could be distinct, with the former EVs being released in a non-lytic fashion through local openings in the cell wall, whereas the latter EVs produced through explosive lysis (**Fig. 4B**).

Taken together, our results draw parallels to bacterial EV biogenesis and suggest that *M. smithii* EVs facilitate the export of DNA fragments covering the entire cellular chromosome, however they are also enriched in eccDNA molecules which originate from excision of a 2.9-kb chromosomal fragment and a proviral genome. Additionally*, M. smithi* EVs may facilitate the export of key metabolic proteins, especially the ones involved in methanogenesis in the gut environment, with potential impact on methane production.

## ACKNOWLEGEMENTS

This work was supported by Agence Nationale de la Recherche (grant ANR-23-CE02-0022 to S.G. and M.K., and grant ANR-22-CE02-0003 to V.C-K.). C.M.G. was supported by an FRM Retour en France fellowship. N.P. was supported by a Pasteur-Roux Postdoctoral Fellowship from the Institut Pasteur (Paris) and the Austrian Science Fond (FWF) Elise Richter Fellowship (FWF project V 931-B). We acknowledge the cryo-ET expertise and assistance of the Institut Pasteur’s NanoImaging Core facility, created and supported by a PIA grant (EquipEx CACSICE: ANR-11-EQPX-0008). We also acknowledge E. Turc and L. Lemée from the Biomics Platform, C2RT, Institut Pasteur, Paris, France, supported by France Génomique (ANR-10-INBS-09) and IBISA. We are grateful for support for Ultrastructural BioImaging Core Facility equipment from the GIS-IBISA, the French Government Programme Investissements d’Avenir France BioImaging (FBI, N° ANR-10-INSB-04-01) and the French government (Agence Nationale de la Recherche) Investissement d’Avenir programme, Laboratoire d’Excellence “Integrative Biology of Emerging Infectious Diseases” (ANR-10-LABX-62-IBEID).

## DATA AVAILABILITY

The mass spectrometry proteomics data have been deposited to the ProteomeXchange Consortium via the PRIDE partner repository with the dataset identifier PXD053033.

## SUPPLEMENTARY MATERIAL

### Material and Methods

#### M. smithii growth conditions

*Methanobrevibacter smithii* PS (ATCC 35061/DSM 861) cultures were grown at 37°C, with agitation at 140 rpm, in serum bottles under strict anaerobic conditions (the gas phase comprised 80% H_2_ and 20% CO_2_ at 2.0 bar) in modified DSM 119 Methanobacterium medium containing 0.5 g/L KH_2_PO_4_, 0.4 g/L MgSO_4_ x 7H_2_O, 0.4 g/L NaCl, 0.4 g/L NH_4_Cl, 0.05 g/L CaCl_2_ x 2H2O, 2 mg/L FeSO4 x 7H2O, 1 mL trace element solution SL-10 (from DSM 320 medium), 1 g/L yeast extract, 1 g/L Na-acetate, 2 g/L Na-formate, 1 mL Selenite-tungstate solution (0.40 g/L NaOH, 6.00 mg/L Na_2_SeO_3_ x 5 H_2_O, 8.00 mg/L Na_2_WO_4_ x 2 H_2_O), 0.5 g/L tryptone, 0.5 mL/L Na-resazurin solution 0.1% w/v, 4 g/L NaHCO_3_, 0.5 g/L L-Cysteine-HCl, 0.5 g/L Na_2_S x 9H_2_O, and 10 mL vitamin solution (from DSM 141 medium). The medium was prepared as described previously (Pende et al., 2021), the pH was adjusted to 7 with HCl.

#### Isolation and purification of EVs

5 mL of an exponentially growing culture of *M. smithii* PS were inoculated into 45 mL of the modified 119 Methanobacterium medium and grown at 37°C as described above. When cultures reached OD_600_ ∼0.35, cells were diluted into fresh modified 119 Methanobacterium medium with an initial OD_600_ of 0.05, grown for 5-8 days and periodically gassed with H_2_ and CO_2_ maintaining the ratio 80:20 until they reach an OD_600_ ∼0.30–0.35. Then, cells were removed by low-speed centrifugation (Eppendorf F-35-6-30 rotor, 7,830 rpm, 20 min, 20°C), supernatant was filtered through 0.45 and 0.22 µm filters (Merck Millipore) and ultracentrifuged to pellet the EVs (120,000 × *g*, 2 h, 15°C, Beckman 45 Ti rotor). After the run, the supernatant was removed, and the pellet was resuspended in 45% Opti-prep^TM^. Different density gradient solutions (20%, 25%, 30%, 35%, 40% and 45% Opti-prep^TM^ with calculated densities 1.110 g/mL, 1.137 g/mL, 1.163 g/mL, 1. 199 g/mL, 1.215 g/mL, and 1.243 g/mL, respectively) were prepared by diluting a 60% Opti-prep^TM^ stock solution (60% Iodixanol; Sigma-Aldrich Chemicals, Zwijndrecht, The Netherlands) with PBS. EVs in 45% Opti-prep^TM^ were layered at the bottom of the ultracentrifuge tube and overlayed with the layers of 40%, 35%, 30%, 25% and 20% solutions. The gradient was ultracentrifuged for 18-24 h at 103,000 ×*g* at 15°C (Beckman SW 60 Ti rotor). After ultracentrifugation, fractions of 0.5 mL were collected from the top of the tube. A portion of each fraction was precipitated with TCA and visualized by SDS-PAGE with the Pierce Silver Stain Kit (Thermo Fisher Scientific) or Coomassie staining according to the manufacturer’s instructions. Fractions were also visualized by TEM. EVs were present in the region of the gradient corresponding to fractions 30-35%. Fractions containing EVs were pooled, diluted in ∼12 mL of PBS and pelleted by ultracentrifugation (220,000 × *g, 3 h,* 15°C, Beckman SW41 rotor). Purified EVs were resuspended in PBS 1X buffer.

#### NanoFCM

The EVs concentration and size distribution was determined by flow cytometry (NanoFCM, Inc., Xiamen, China), as previously described (Grasekamp et al., 2023). The instrument was aligned, focused, and calibrated using 0.25 μm Fluorescent Silica Microspheres, that also served as concentration standard. Silica Nanosphere Cocktail (S16M-Exo) was used as a size standard (NanoFCM, Inc.; range 68-155nm), from which a calibration curve was calculated and used to infer the size of the events present in each sample using the NanoFCM software (NanoFCM Profession V2.0). All samples were diluted with 0.02 µm filtered PBS 1X solution to ensure the particle count was within the range of 2000–12,000/min. All particles that passed by the detector over 60-second intervals were recorded.

#### Transmission electron microscopy (TEM)

Before staining, carbon-coated copper grids were incubated with a 0.1% of poly*-*L*-*lysine solution in H_2_O for 30 min (Barth, 1985). *Coated grids were washed three times with water and subsequently* 5 µL of the sample was applied to the grid and negatively stained using 2% uranyl acetate (wt/vol). Samples were imaged with the transmission electron microscope FEI Spirit Tecnai Biotwin operating at 120 kV. The relative diameter of EVs was determined as previously described (Liu et al., 2021). Briefly, electron micrographs of negatively stained EVs were analyzed with ImageJ (Rueden et al., 2017), and the area of each was calculated manually (n= 412). The relative diameter was calculated according to the equation A=(π/4) × D^2^

#### DNA isolation from EVs and sequencing

Before DNA extraction, EVs were treated with DNase I and RNase in the presence of MgCl_2_ to remove cellular DNA, as described before (Liu et al., 2021). Subsequently, EVs were disrupted by incubation with SDS and proteinase K at final concentrations of 0.5% (wt/vol) and 100 µg/ml, respectively, for 30 min at 55°C. The DNA was extracted with the mixture of phenol/chloroform/isoamyl alcohol (25:24:1 vol/vol/vol) and precipitated with sodium acetate to a final concentration of 0.3 M and ice-cold 70% ethanol. Sequencing libraries were prepared and sequenced on Illumina MiSeq platform with 150-bp paired-end read lengths (Institut Pasteur, France). Raw sequence reads were processed with Trimmomatic v.0.3.6 (Bolger et al., 2014) and mapped to the reference genome of *M. smithii* PS using Bowtie2 with default parameters (Langmead et al., 2009) and visualized with UGENE (Okonechnikov et al., 2012). In addition, raw sequences were assembled with MetaSPAdes v3.11.1 with default parameters (Nurk et al., 2017). For the contig corresponding to the extrachromosomal circular element, open reading frames (ORFs) were predicted by Prokka v.1.14.5 (Seemann, 2014). Searches for distant homologs were performed using HHpred against PFAM, PDB and CDD databases (Söding et al., 2005).

#### Mass spectrometry and data analysis

The protein content of *M. smithii* PS cells and purified EVs (triplicates) were analyzed by liquid chromatography – tandem mass spectrometry (LC-MS/MS) at the Proteomics Platform of Institut Pasteur (Paris, France) as previously described (Liu et al., 2021). Samples were snap-frozen in liquid nitrogen, lyophilized and re-suspended in 100 μl of lysis buffer including 8 M Guanidine HCl (GuHCl), 5 mM Tris(2-carboxyethyl)phosphine (TCEP). Samples were sonicated in a Covaris E220 (Covaris) for 5 min at 200 cycles/burst, with 175 W peak power and a 10% duty cycle. Lysates were centrifuged 15 min, 15,000 g at RT to and supernatants were kept. 2-chloro-acetamide (CAA) was added to a final concentration of 20 mM. Subsequently, samples were incubated at 95°C for 5 min, and 9 times volume samples of 50 mM Tris-HCl (pH 8.0) were added to dilute GuHCl to a concentration of under 1M. A mixture of 2 µg of Trypsin/Lys-C (Promega – V5071) was added to the samples and kept at 37°C overnight for digestion of the proteins. The reaction was stopped by addition of formic acid (FA) at 1% final. Peptides were desalted using Sep-Pac C18 Cartridges (Waters, USA) and eluted with 80% ACN / 0.1% FA. The purified peptides were concentrated to near dryness, re-suspended in 25 μl of 2% ACN / 0.1 % FA and analyzed by Nano LC-MS/MS using an EASY-nLC 1200 system (peptides were loaded and separated on a 50 cm long home-made C18 column; 75 µm ID, 1.9 µm particles, 100 Å pore size, ReproSil-Pur Basic C18 - Dr. Maisch GmbH, Ammerbuch-Entringen, Germany, coupled to an Orbitrap Lumos tribrid (Thermo Fisher Scientific) tuned to the DDA mode). Peptides were eluted with a multi-step gradient from 5 to 25% buffer B (ACN 80% / FA 0.1%) in 95 min, 25 to 40% buffer B in 15 min and 40 to 95% Buffer B in 10 min at a flow rate of 250 nL/min for up to 130 min. Column temperature was set to 60°C.

Mass spectra were acquired using Xcalibur software using a data-dependent Top 2s method with a survey scans (300-1700 m/z) at a resolution of 60,000 and MS/MS scans (fixed first mass 110 m/z) at a resolution of 15,000. The AGC target and maximum injection time for the survey scans and the MS/MS scans were set to 6.0E+05, 50ms and 5.0E+04, 100ms respectively. The isolation window was set to 1.6 m/z and normalized collision energy fixed to 30 for HCD fragmentation. We used a minimum intensity threshold of 5E+04. Precursor ion charge states from 2 to 7 were accepted and advanced peak determination was enabled. Exclude isotopes was enabled and selected ions were dynamically excluded for 45 seconds.

Peptide masses were searched against a UniProt *M. smithii PS* database (1,783 entries the 19/01/2023) using Andromeda (Cox et al., 2011) with the MaxQuant ver. 2.0.3.0 (Tyanova et al., 2016) software. Variable modifications (methionine oxidation and N-terminal acetylation) and fixed modification (cysteine carbamidomethylation) were set for the search and trypsin with a maximum of two missed cleavages was chosen for searching. The minimum peptide length was set to 7 amino acids and the false discovery rate (FDR) for peptide and protein identification was set to 0.01. The main search peptide tolerance was set to 4.5 ppm and to 20 ppm for the MS/MS match tolerance. Second peptides were enabled to identify co-fragmentation events. Identified proteins were functionally annotated using the archaeal clusters of orthologous groups (arCOG) database (Makarova et al., 2015). Differential expression of proteins was calculated using the R package DEP (differential enrichment analysis of proteomics data) (v. 1.21.0) as described in (Mills et al., 2024; Zhang et al., 2018). Data underwent variance stabilizing transformation for normalization with the vsn function in the DEP package. The threshold for significant enrichment in EVs compared to cells is log_2_ fold change greater than 1.5 and adjusted *P*-value lower than 0.05.

The mass spectrometry proteomics data have been deposited to the ProteomeXchange Consortium via the PRIDE (Perez-Riverol et al., 2022) partner repository with the dataset identifier PXD05303.

#### Detection of the extrachromosomal circular element by PCR

Polymerase chain reactions (PCRs) with primers targeting the integrated (F: 5’-TCTTCAGGACTTACATCCAGG-3’, R: 5’-TGTACGTTCACATCCGTCTA-3’; expected size: 291 bp) and excised (F: 5’-CTGTTGAAGAAGGTAAACCCG-3’, R: 5’-TTGTACGTTCACATCCGTCT-3’; expected size: 433 bp) form of the extrachromosomal circular element were performed both on *M. smithii* cells and purified EVs. PCRs were performed using DreamTaq DNA Polymerase with the following steps: Denaturation at 95°C for 3 min followed by 33 cycles of 95°C 30 s, 57°C 30 s, 72°C 1 min, and a final extension step at 72°C for 10 min. The sequences of the amplified products were confirmed by Sanger sequencing and analyzed with QIAGEN CLC Main Workbench 24.0 (QIAGEN, Aarhus, Denmark).

#### Detection of MSTV1 by PCR

PCRs with primers targeting the integrated (F: 5’-GGGTTTAATTTTGGGGGATA-3’, R: 5’-AGGATTTCTTCATTGGTTCTCA-3’) and excised (F: 5’-TTGATGATGTTAATAATGGTGATGA-3’, R: 5’-AGGATTTCTTCATTGGTTCTTCTCA-3’) forms of MSTV1 were performed both on *M. smithii* cells and purified EVs (Baquero et al., under review). PCRs were performed using DreamTaq DNA Polymerase with the following steps: denaturation at 95°C for 3 min followed by 35 cycles of 95°C 30 s, 57°C 30 s, 72°C 1 min, and a final extension step at 72°C for 10 min.

#### Cryo-ET: Sample preparation and tilt series acquisition

Samples for cryo-electron tomography were prepared as described previously (Baquero et al., under review). Briefly, a solution of bovine serum albumin–gold tracer containing 10-nm-diameter colloidal gold particles was added to a fresh culture of *M. smithii* in its exponential phase with a final ratio of 1:1. A small amount of the sample was applied to the glow discharged (ELMO, Corduan) carbon-coated copper grids (Cu 200 mesh Quantifoil R2/2). The sample was rapidly frozen in liquid ethane using a Leica EMGP system. The grids were stored in liquid nitrogen until image acquisition. Tilt series were collected on a 300 kV Titan Krios G3 transmission electron microscope (Thermo Fisher Scientific) equipped with a X-FEG Tip, a Gatan K3 Direct Electron Detector and a Gatan BioQuantum LS Imaging Filter with slit width of 20 eV and a single-tilt axis holder. Tilt series were acquired with Tomography software v.5.6 (Thermo Fisher Scientific) using a dose-symmetric scheme (Hagen et al., 2017), with an angular range of ±60°, 2° angular increment, -8 µm defocus, pixel size of 3.4 Å (26000x). The total dose was set at 140 e/Å² at a dose rate of 41.6 e/pix/sec in vacuum, with C2 and Objectif Apertures of 100 µm. 3D tomographic reconstructions were calculated in IMOD by weighted back projection using the SIRT-like filter with 9 iterations (Mastronarde and Held, 2017).

#### Cryo-ET: Segmentation and analysis of tomographic data

The drawing tools of IMOD (Mastronarde and Held, 2017) were used for tomogram annotation. Archaeal membrane and cell wall were manually traced every 30 slides and the subsequent use of the interpolator tool. Both closed and open contours were employed, depending if the full cell was displayed in the field of view. EVs were modeled by manual tracing in all the slices where they were present. All traces were merged through the “merge” tool. Surfaces were generated using the ‘imodmesh’ function. The IMOD surfaces were then imported to UCSF ChimeraX (Meng et al., 2023) together with the tomographic file. IMOD surfaces were either used directly in visualization or used to mask out regions of the tomographic volume with the ‘volume mask’ tool of ChimeraX. The subtomographic areas of interest were then visualized in iso-surface representations of variant threshold values and colors.

---

**Figure S1.**
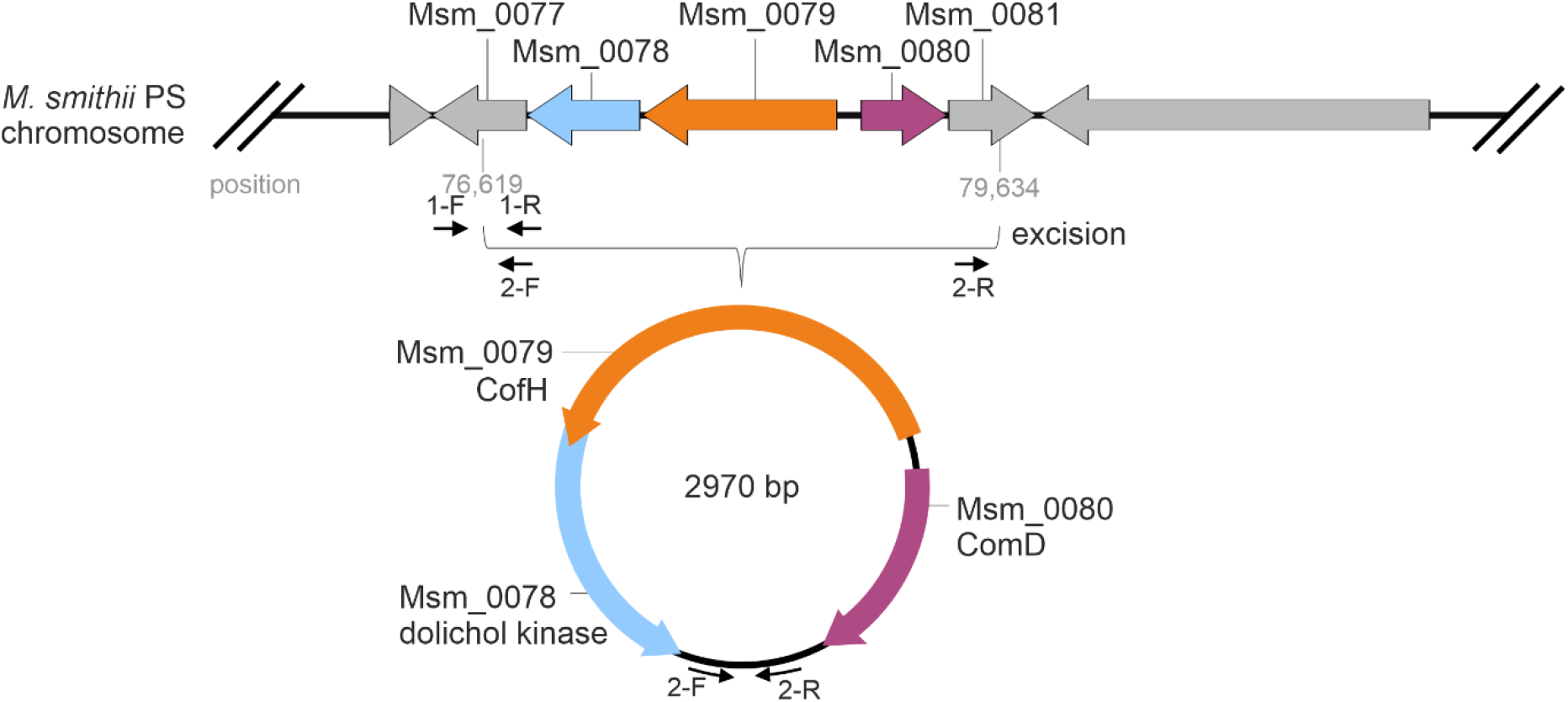
Schematic representation of the excision of the 2970 bp circular element. Two sets of primers (indicated with small black arrows) were designed to detect the integrated and excised forms of the virus genome. The primers 1-F and 1-R targets the sequence integrated into the host chromosome (1-F 5’ TCTTCAGGACTTACATCCAGG, R 5’ TGTACGTTCACATCCGTCTA). Primers 2-F and 2-R detects the circularized form of the extrachromosomal element (2-F: 5’-CTGTTGAAGAAGGTAAACCCG-3’, 2-R: 5’-TTGTACGTTCACATCCGTCT-3’). The PCR results are shown in Fig. 2C.

**Figure S2.**
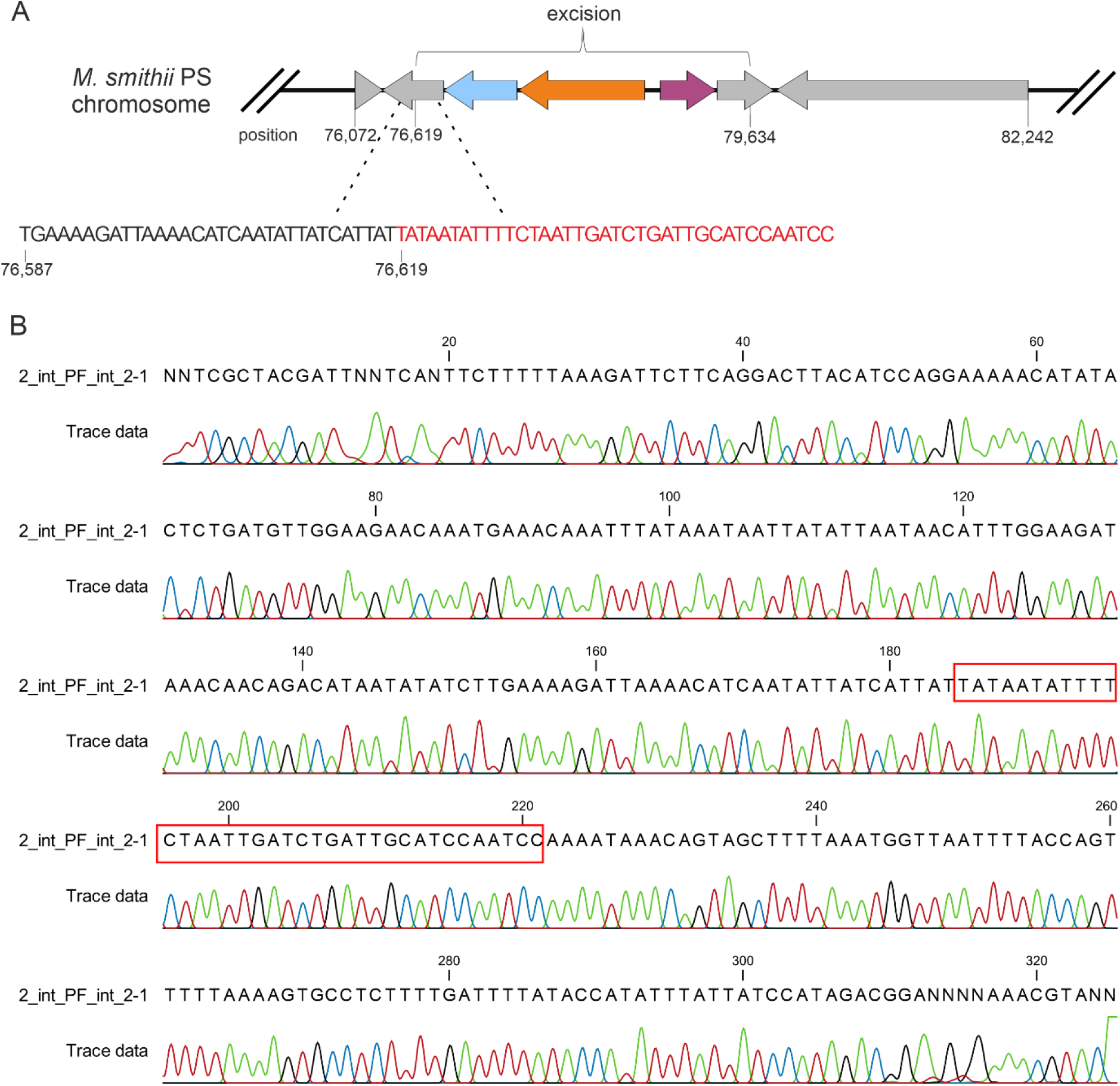
Detection of the element integrated into the host chromosome. A, Schematic representation showing the sequence of the circular element integrated into the host genome. B, Sequencing chromatogram confirming the results obtained by PCR (Fig. 2C).

**Figure S3.**
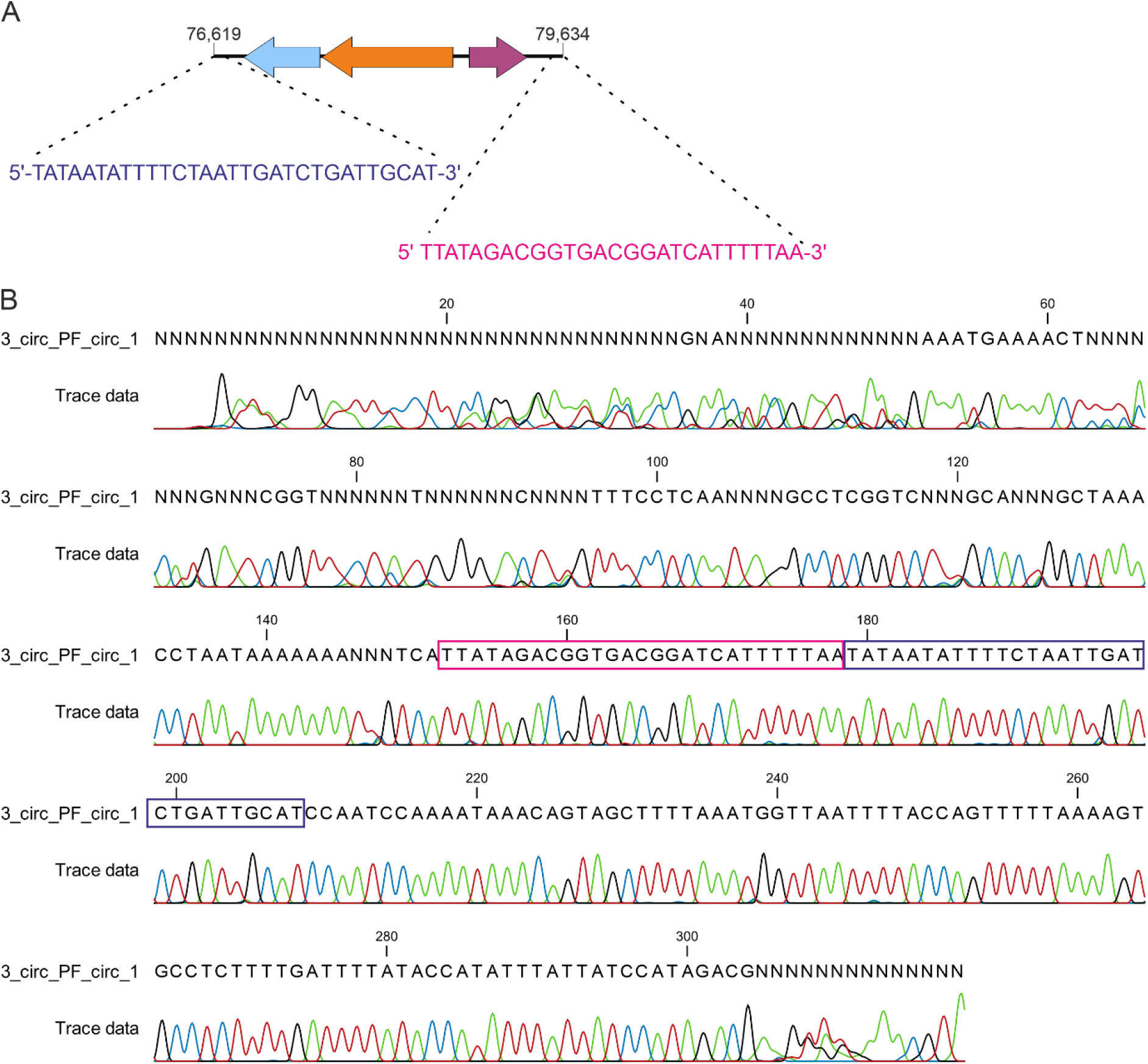
Detection of the circularized extrachromosomal element in *M. smithii* EVs. A, Schematic representation of the sequences at both termini of the element. B, Sequencing chromatogram confirming the results obtained by PCR (Fig. 2C).

**Figure S4.**
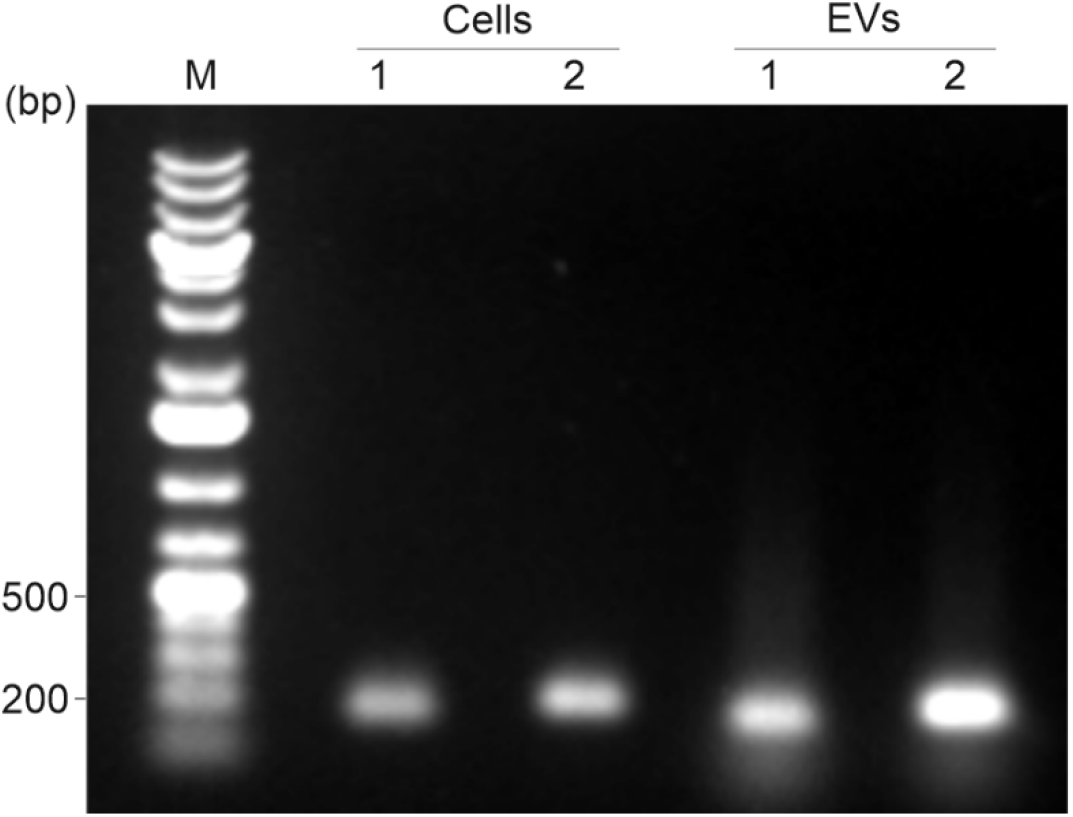
Detection of MSTV1 in both cells and purified EVs of *M. smithii* PS. The agarose gel electrophoresis shows the amplified products: lane 1, provirus integrated into the host chromosome (F: 5’-GGGTTTAATTTTGGGGGATA-3’, R: 5’-AGGATTTCTTCATTGGTTCTCA-3’; expected size: 180 bp); lane 2, excised and circularized form of the MSTV1 genome (F: 5’TTGATGATGTTAATAATGGTGATGA-3’, R: 5’-AGGATTTCTTCATTGGTTCTTCTCA-3’; expected size: 216 bp). MW: Thermo Scientific GeneRuler 1 kb Plus DNA Ladder.

**Figure S5.**
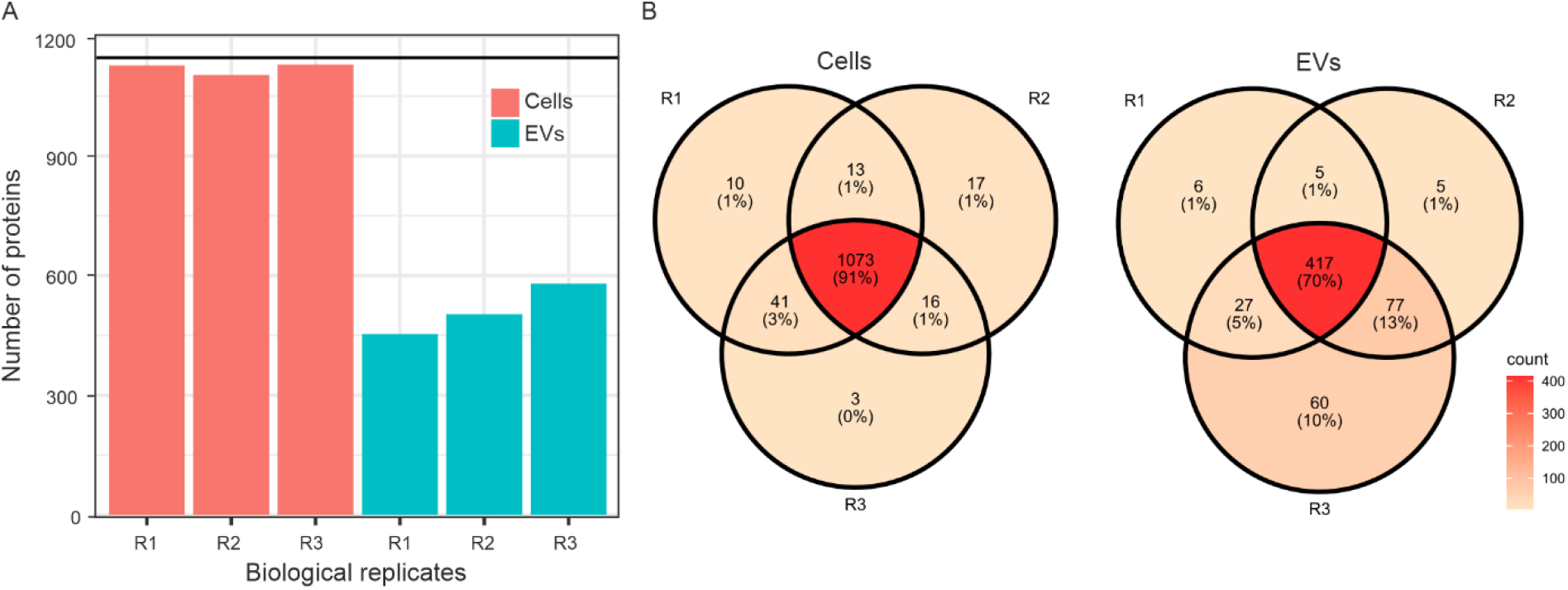
Proteomic analysis of *M. smithii* cells and EVs. A, Proteins identified per replicate for *M. smithii* cells and EVs. B, Venn diagram of the identified proteins per replicate for cells and EVs. The number and percentage reported in each circle indicate the total and relative amount of proteins found per sample.

